# Neuroectoderm phenotypes in a human stem cell model of O-GlcNAc transferase intellectual disability

**DOI:** 10.1101/2023.09.18.558285

**Authors:** Marta Murray, Lindsay Davidson, Andrew T. Ferenbach, Dirk Lefeber, Daan M. F. van Aalten

## Abstract

Most intellectual disabilities are caused by monogenic variation. Mutations in the O-GlcNAc transferase (*OGT*) gene have recently been linked to a novel congenital disorder of glycosylation (OGT-CDG), involving symptoms of possible neuroectodermal origin. To test the hypothesis that pathology is linked to defects in differentiation during early embryogenesis, we developed an OGT-CDG induced pluripotent stem cell lines together with isogenic controls generated by CRISPR/Cas9 gene-editing. Although the OGT-CDG variant leads to a significant decrease in OGT and O-GlcNAcase protein levels, there were no changes in differentiation potential or stemness. However, differentiation into ectoderm resulted in significant differences in O-GlcNAc homeostasis. Further differentiation to neuronal stem cells revealed differences in morphology between patient and control lines, accompanied by disruption of the O-GlcNAc pathway. This suggests a critical role for O-GlcNAcylation in early neuroectoderm architecture, with robust compensatory mechanisms in the earliest stages of stem cell differentiation.

## Introduction

Intellectual disability (ID) affects 1-3% of the human population and is caused by environmental and genetic factors, the latter including chromosomal rearrangements, gene duplications and single nucleotide polymorphisms. Over 160 rare genes have been identified as associated with glycosylation (<u>Varki, 2022</u>) Recently, a novel congenital disorder of glycosylation was identified, arising from variants in O-GlcNAc transferase (OGT), named OGT-CDG (<u>Pravata et al., 2019</u>, <u>Pravata et al., 2020b</u>, <u>Niranjan et al., 2015</u>). OGT modifies serine and threonine residues of target proteins with the sugar moiety GlcNAc (<u>Haltiwanger</u> <u>et al., 1990</u>, <u>Hanover et al., 2003</u>). This post-translational modification is reversible, and hydrolysis of the sugar from target proteins is mediated by O-GlcNAc hydrolase (OGA) (<u>Dong and Hart, 1994</u>). O-GlcNAcylation critically affects protein localization, stability and activity, and potentially regulates other post-translational modifications (<u>Golks et al., 2007</u>, <u>Lo et al., 2018</u>, <u>Ruan et al., 2013</u>). Globally, O-GlcNAcylation affects cell cycle, translational and epigenetic cellular regulation (<u>Yang and Qian, 2017</u>, <u>Jackson and Tjian,</u> <u>1988</u>, <u>Liu and Li, 2018</u>). Unlike phosphorylation, O-GlcNAcylation affects over 5000 target proteins utilizing only a single pair of enzymes that are ubiquitously expressed (<u>Haltiwanger et al., 1990</u>, <u>Dong and Hart,</u> <u>1994</u>). For example, neuronal connectivity is critically dependent on O-GlcNAcylated proteins, both for synapse formation and function (<u>Lagerlof et al., 2017</u>, <u>Wheatley et al., 2019</u>). Feedback regulation controls both OGA and OGT physiological levels and activity (<u>Lin et al., 2021</u>, <u>Muha et al., 2021</u>, <u>Krzeslak et al.,</u> <u>2012</u>, <u>Forster et al., 2014</u>), although the mechanisms underpinning this are not yet understood. Pathogenic variants in OGT-CDG impact this feedback and overall O-GlcNAcylation levels (<u>Forster et al., 2014</u>, <u>Pravata</u> <u>et al., 2019</u>, <u>Omelkova et al., 2023</u>).

OGT consists of a C-terminal catalytic domain, and an N-terminal tetratricopeptide repeat (TPR) domain (<u>Iyer and Hart, 2003</u>) suggested to modulate substrate specificity and binding (<u>Iyer and Hart, 2003</u>, <u>Burrows et al., 2016</u>, <u>Rafie et al., 2017</u>, <u>Stephen et al., 2021</u>). Different tissues have different levels of OGT and OGA, leading to different levels of O-GlcNAcylation in terminally differentiated cells but also during development (<u>Shafi et al., 2000</u>, <u>Muha et al., 2021</u>, <u>Cheng et al., 2020</u>). Moreover, disbalances in the substrate O-GlcNAcylation levels and changes in O-GlcNAc homeostasis are associated with neurodegenerative disease, cancer, diabetes, and cardiovascular diseases (<u>Lee et al., 2021b</u>, <u>Zhu and Hart,</u> <u>2021</u>, <u>Nie and Yi, 2019</u>). It is unclear what mechanisms link mutations in OGT to OGT-CDG symptoms (<u>Pravata et al., 2020b</u>). It is also unknown whether the effects are mainly induced through, for instance, changes in OGT levels or changes in OGT activity. Therefore, a critical assessment of O-GlcNAcylation homeostasis in models of OGT-CDG is required to understand the etiology of this disease and potential tissue-specific mechanisms for regulation (<u>Shafi et al., 2000</u>, <u>Parween et al., 2017</u>).

Analysis of the O-GlcNAc pathway of several pathogenic OGT variants modeled in different systems has revealed a range of symptoms (<u>Pravata et al., 2019</u>, <u>Willems et al., 2017</u>, <u>Pravata et al., 2020a</u>, <u>Vaidyanathan et al., 2017</u>, <u>Bouazzi et al., 2015</u>, <u>Selvan et al., 2018</u>, <u>Fenckova et al., 2022</u>). Pathogenic mutations occur in both domains, affecting stability and/or activity, which consequently result in effects on O-GlcNAc homeostasis (<u>Pravata et al., 2020b</u>). Patients carrying pathogenic OGT variants commonly show moderate to severe intellectual disability accompanied with developmental delay, psychomotor retardation, and behavioral problems. Concomitantly, several patients display facial dysmorphias (<u>Pravata et al., 2020b</u>). Despite an ability to group mutations according to variant location and their effects on catalytic glycosyltransferase activity, extracting generalizations of the current literature is challenging. An additional difficulty in systematic comparison arises when comparing effects observed across different model systems, ranging from embryonic stem cells to fly. These approaches differ in key points like different genetic backgrounds (<u>Cooper et al., 2013</u>), metabolism (<u>Jozwiak et al., 2014</u>), proliferation rates (<u>Levine et al.,</u> <u>2021</u>), and both capacity and acute requirement for O-GlcNAcylation (<u>Vaidyanathan and Wells, 2014</u>, <u>Shafi</u> <u>et al., 2000</u>, <u>Omelkova et al., 2023</u>). Therefore, the effects of OGT-CDG mutants on specific tissues and on developmental processes such as differentiation are not yet known.

The O-GlcNAc pathway significantly impacts both embryonic stem cells and neurons (<u>Andres et al., 2017</u>, <u>Kim et al., 2021</u>, <u>Myers et al., 2016</u>, <u>Wang et al., 2016</u>, <u>Su and Schwarz, 2017</u>, <u>Chen et al., 2021</u>). O-GlcNAcylation is essential for neuronal function. Moreover, it is clear that the O-GlcNAc pathway has a continual role in brain development (<u>Liu et al., 2012</u>). However, the early stages of neural development leading from epiblast formation to neuroepithelium remain largely unexplored. Early genetic programs result in formation of the epiblast (at embryonic day 3.5 in mouse, day 5 in human) (<u>Rossant and Tam, 2022</u>). This cell mass consists of totipotent stem cells, differentiating to all organismal tissues. Following epiblast formation, gastrulation results in formation of the three germ layers (ectoderm, endoderm and mesoderm; mouse E6-E9.5, human E14-16) (<u>Muhr and Ackerman, 2023</u>). These three layers subsequently and distinctly give rise to all further organs and tissues, except for trophoblast. During this stage, stresses and insults from gene and protein imbalance result in abnormal development (<u>Pini et al., 2020</u>, <u>McMillan et al.,</u> <u>2018</u>). Intriguingly, many of the symptoms displayed by patients harboring OGT-CDG variants could mainly arise from ectoderm-derived tissues. Therefore, we hypothesize that ectoderm differentiation could be altered in cells carrying pathogenic variants, and more broadly, disbalance in O-GlcNAcylation will impact on early development.

Here, we modeled an OGT variant in the cellular context of early human neuroectoderm differentiation using patient-derived induced pluripotent stem cells (iPSCs). Although there are no changes in the OGT-CDG variant in terms of global O-GlcNAcylation in undifferentiated iPSCs, there are significant decreases in OGT and OGA protein levels. Further differentiation to neural rosettes induced reduction of O-GlcNAc levels accompanied by similar decreases in OGT and OGA protein levels, suggesting that changes in compensatory mechanism, as compared to iPSCs, are critical in regulating total O-GlcNAcome. A phenotypic investigation into the morphology revealed that neural rosettes generated from the OGT-CDG iPSCs were smaller and had organization defects. Overall, modeling tissue specific regulation of O-GlcNAcylation demonstrates a striking differential regulation of the OGT-OGA balance, shifting from transcriptional to protein-dependent.

## Results and Discussion

### Correction of an OGT-CDG variant restores normal splicing in IPSCs

To model OGT-CDG and its effects on O-GlcNAc homeostasis, we focused on a previously reported pathogenic variant in *OGT* harboring a single nucleotide change located in the intron-exon junction of intron 3 (Fig. 1A). This variant results in skipping of exon 4 from the majority of transcripts (<u>Willems et al., 2017</u>). In patient fibroblasts, this variant has decreased levels of OGT and OGA with no apparent change in O-GlcNAcylation (<u>Willems et al., 2017</u>).

**Figure 1:**
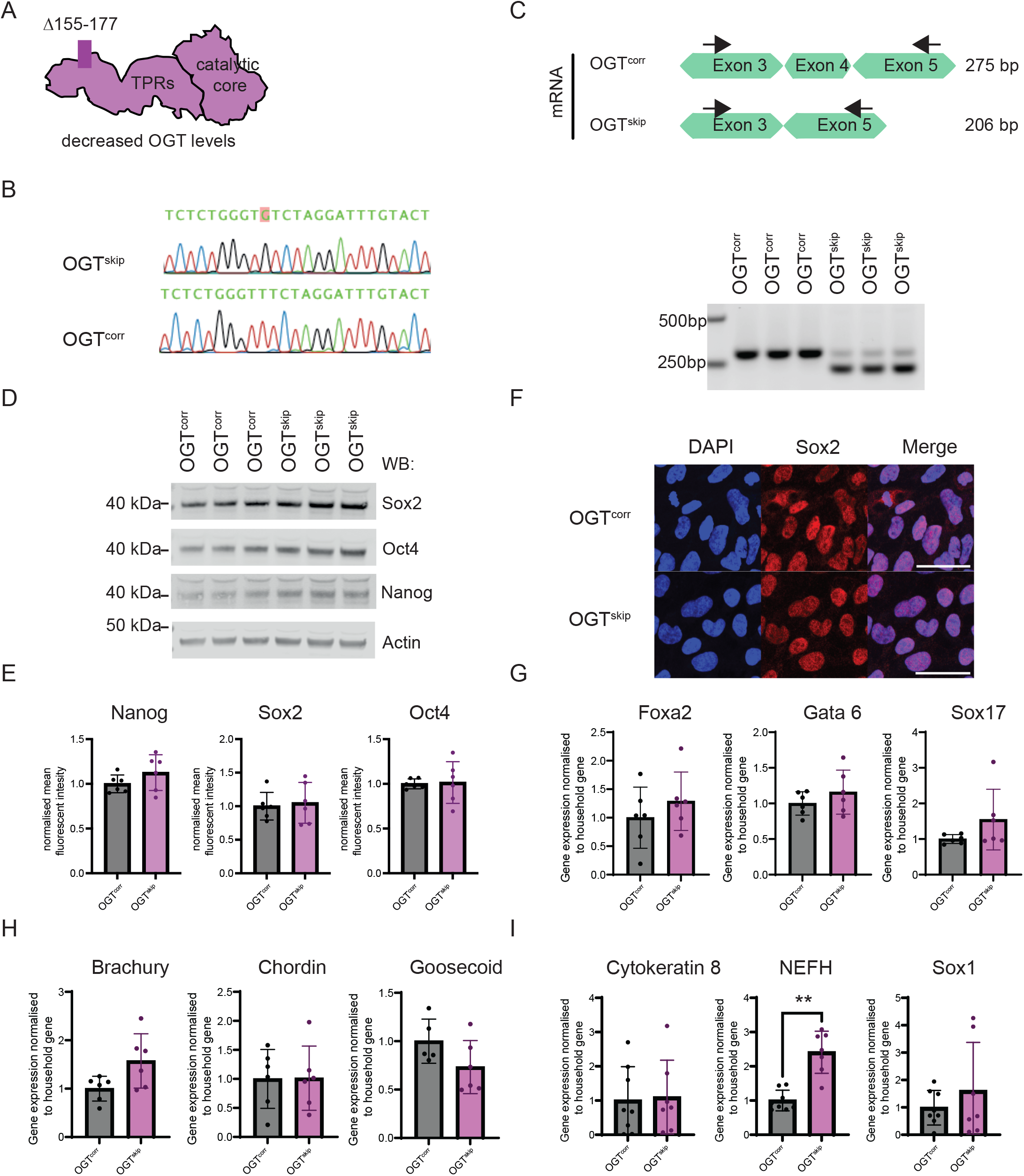
Generation of corrected genome-matched IPSCs. **(a)** Schematic representation of OGT structure consisting of the TPRs and catalytic domain. The mis-splicing of the patient variant leads to a protein product that lacks amino acids 155-170. **(b)** Representative DNA sequencing chromatograms from the genomic region of the corrected nucleotide. **(c)** Schematic and PCR of mRNA isolated from corrected and patient IPSCs. Schematic demonstrates theoretical PCR product size in the variants. Agarose gel demonstrates presence of longer transcript in the correct IPSC cell lines while the patient derived cell predominantly shows the misspliced transcript. **(d)** Representative image and **(e)** quantification of Western blot for Yamanaka factors in three clonal lines demonstrating no changes to stem cell markers. N = 2 experiments, 3 clonal lines/biological replicates. **(f)** Representative confocal microscopy image of DAPI and Sox2 immunofluorescence depicting no changes to stem cell markers at single cell resolution. Scale bar, 50 μm. **(g)** Differentiation potential by qPCR of specific markers in the OGT^corr^ and OGT^skip^ cell lines for endoderm, **(h)** Mesoderm, and **(i)** Ectoderm. n = 2 experiments, 3 clonal lines/biological replicates for each **(g)-(i)**.

Patient derived IPSCs harboring the exon 4 skipping variant (referred to as OGT^skip^) were generated from the male proband. Genetic background heterogeneity is a large source of variability in both IPSCs (<u>Burrows</u> <u>et al., 2016</u>, <u>Kilpinen et al., 2017</u>, <u>Kyttala et al., 2016</u>) and patient symptoms (<u>Cooper et al., 2013</u>). Using CRISPR/Cas9-mediated homologous recombination, three genome-matched OGT wild type (OGT^corr^) cell lines were created, differing from the patient line only in the pathogenic variant. To verify nucleotide substitution at the targeted site, Sanger sequencing of genomic DNA was employed (Fig. 1B). This demonstrated the correct non-homologous recombination as designed, with no coding changes in the direct vicinity of the target site. To ensure the sequence variation in OGT^corr^ and OGT^skip^ exists only in the desired position, the entire mRNA open reading frame was sequenced. This confirmed successful correction of the single targeted codon and the presence of silent mutations of the gRNA. From the Cas9-treated cell pool, we also selected three unedited clones, which were used as the patient variant in all subsequent experiments, designated OGT^skip^ from hereon. The transcript product size was evaluated by RT-PCR (Fig. 1C), and in agreement with an initial report on this variant (<u>Willems et al., 2017</u>), OGT^skip^ predominantly gives rise to shortened transcripts, with a minimal portion of full-length transcript attributable to splicing escape – and presumably survival of the patient. In contrast, the OGT^corr^ lines were confirmed to produce only full-length transcripts (Fig. 1C). Taken together, these experiments have generated genome-matched lines differing only in a single site, yielding an IPSCs model of OGT-CDG together with a corrected control that restores normal splicing.

### OGT^skip^ and OGT^corr^ IPSCs retain the hallmarks of stemness

As OGT influences Sox2 and Oct4 activity in human ESCs (<u>Myers et al., 2016</u>, <u>Kim et al., 2021</u>, <u>Jang et al.,</u> <u>2012</u>, <u>Constable et al., 2017</u>) we evaluated levels of the central Yamanaka factors that maintain cell stemness (<u>Takahashi and Yamanaka, 2006</u>). No differences in the overall levels of the stem cell markers nanog, Oct4, and Sox2 were observed between the OGT^skip^ and OGT^corr^ iPSCs as evaluated by Western blot (Fig. 1D, E) and qPCR (Supplementary Fig. 1A). To assess levels of these stem cell markers and effects on morphology at the single cell level, we examined the same key factors by immunofluorescence (Fig. 1F, and Supplementary Fig. 1B-D) and flow cytometry (Supplementary Fig. 1E), revealing no differences between the OGT^skip^ and OGT^corr^ lines. To ensure cells were pluripotent, they were passaged to assure self-renewal, and differentiated into the three germ layers. Differentiation was assessed by qPCR, showing that both OGT^corr^ and OGT^skip^ differentiated successfully without differences (Fig. 1G-I).

All hallmarks of stemness were equivalent in patient and control variant lines (Fig. 1 and Supplementary Fig. 1). Therefore, these results provide direct evidence that all cell lines have retained their characteristics as stem cells, with no detectable disruption of stemness and pluripotency.

### OGT^skip^ IPSCs maintain global O-GlcNAcylation levels by downregulating OGA

IPSCs are biochemically and functionally representative of the inner cell mass of the pre-implantation blastocyst (<u>Takahashi et al., 2007</u>, <u>Takahashi and Yamanaka, 2006</u>). To assess the O-GlcNAc pathway in IPSCs, we measured levels of OGT, OGA, and total O-GlcNAc (Fig. 2A, B). OGT and OGA levels in the OGT^skip^ cell line were both decreased to 57% of OGT^corr^. These decreases were not accompanied by changes in overall O-GlcNAcylation. To determine whether the observed changes were transcriptionally derived, we assessed levels of *OGT* and *OGA* transcripts by qPCR. The levels of *OGT* transcripts revealed an increase (22%) in *OGT* mRNA (Fig. 2C), suggesting that increasing transcript levels is used as a mechanism to compensate for missplicing. Moreover, we observe a significant decrease (to 51%) in *OGA* transcript levels. Taken together, these data suggest an intricate balance in the O-GlcNAc pathway in IPSCs. The pathogenic OGT variant results in decreased OGT protein levels, accompanied by decreased OGA levels. This response, which likely arises from transcription, provides IPSCs with buffering for total O-GlcNAc allowing OGT^skip^ IPSCs to maintain physiological O-GlcNAcome levels through downregulation of OGA.

**Figure 2:**
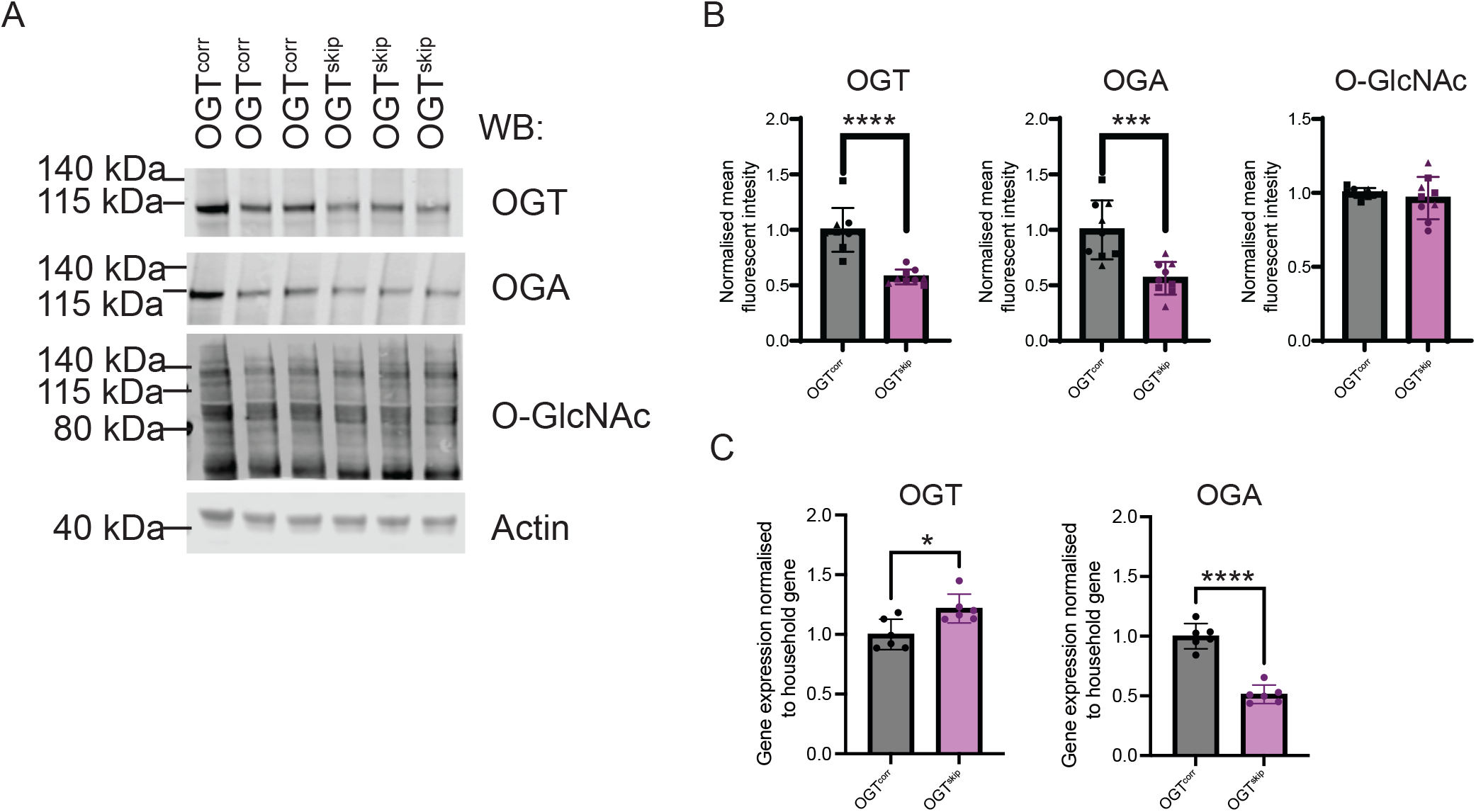
OGT^SKIP^ IPSCs maintain protein O-GlcNAcylation levels through downregulation of OGA. **(a)** Representative Western blot of IPSCs and **(b)** Quantification of changes in OGT, OGA, and total O-GlcNAc. n = 3 experiments, 3 clonal lines/biological replicates for each. **(c)** Changes in transcription levels by qPCR.

### Ectoderm differentiation reveals decreased protein O-GlcNAcylation

Differentiation of pluripotent cells into neurons is a stepwise process, leading from stem cells to epiblast/ectoderm, then neuroectoderm/neural progenitors, and ultimately a defined neuronal cell fate. OGT inhibition leads to accelerated differentiation of human embryonic stem cells to neurons (<u>Andres et al., 2017</u>), suggesting that differentiation potential is directly influenced by O-GlcNAcylation (<u>Pravata et al., 2019</u>, <u>Omelkova et al., 2023</u>). Moreover, conditional knock-down of OGT in neurons leads to the loss of neuronal stem cells in mice (<u>Chen et al., 2021</u>). However, it is unclear at which point during the differentiation pathway changes in O-GlcNAc homeostasis are critical for survival. To begin to address this, we first differentiated IPSCs into ectoderm, (Figure 1I and Figure 3). Differentiation efficiency was defined by qPCR using markers such as Sox1, NEFH, and cytokeratin 8. Apart from a significant increase in levels of NEFH, a neurofilament and a marker for ectoderm, all other ectoderm-specific markers analyzed were unchanged in OGT^skip^ compared to the OGT^corr^ lines, suggesting equal levels of differentiation into ectoderm.

**Figure 3:**
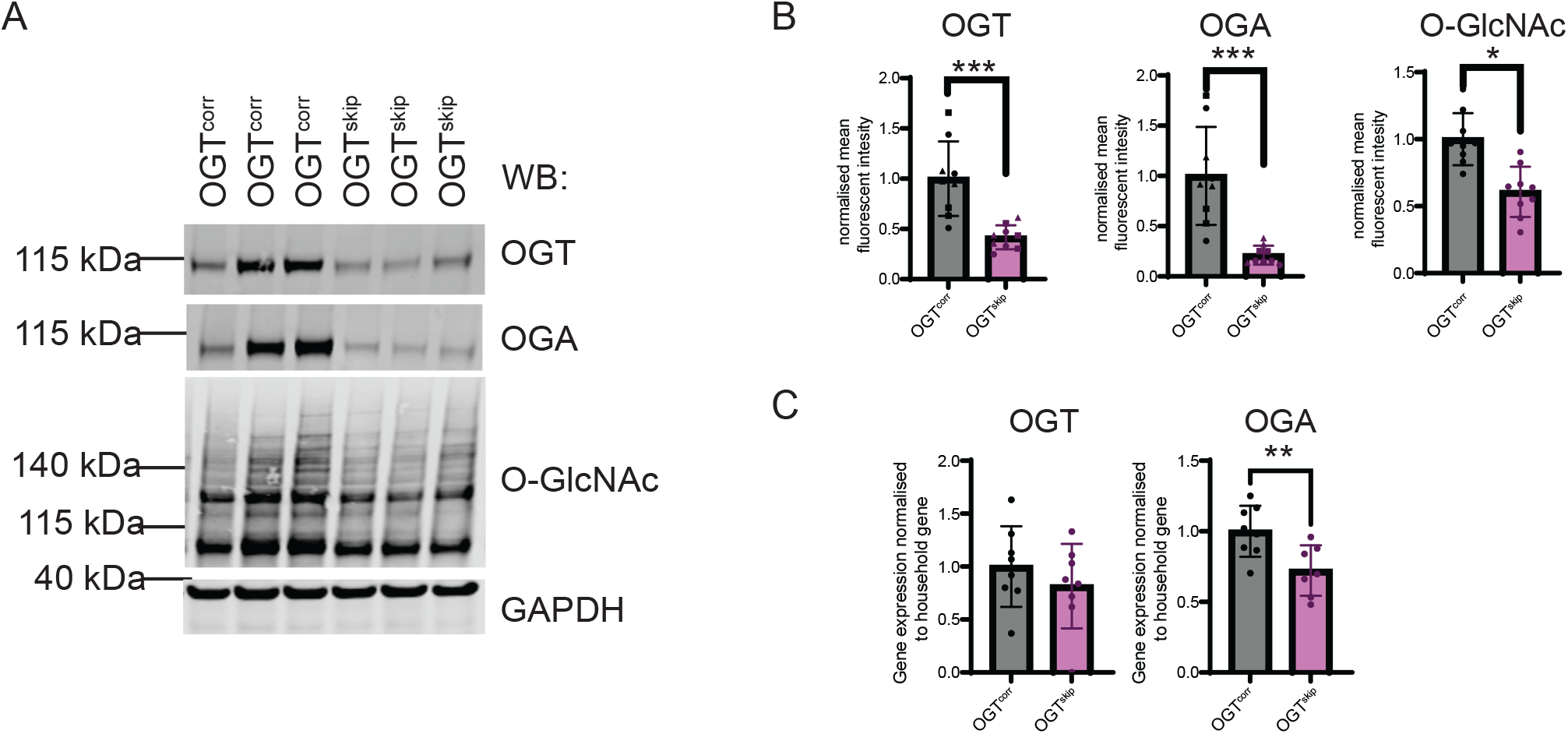
OGT^SKIP^ ectoderm reveals decreased levels of protein O-GlcNAcylation. **(a)** Representative Western blot of ectoderm-differentiated IPSCs and **(b)** Quantification of changes in OGT, OGA, and total O-GlcNAc. n = 3 experiments, 3 clonal lines/biological replicates for each. **(c)** Changes in transcription levels of OGT and OGA by qPCR.

Comparison of the OGT, OGA, and O-GlcNAc levels between OGT^skip^ and OGT^corr^ IPSCs differentiated towards ectoderm revealed a significant decrease of OGT protein levels (to 42%) and a more pronounced decrease of OGA protein levels (to 21%) in OGT^skip^ ectoderm (Fig. 3A, B). This was accompanied by a significant decrease in the total O-GlcNAcome in OGT^skip^ to 61% of OGT^corr^. To assess whether these changes were of transcriptional origin, we measured mRNA levels (Fig. 3C), revealing no detectable differences in *OGT* transcripts between the OGT^skip^ and OGT^corr^ IPSCs. In contrast, *OGA* transcript levels in the OGT^skip^ variant were significantly decreased (to 72%) compared to OGT^corr^, suggesting a shift in OGA protein regulation from transcriptional in IPSCs to post-transcriptional in the ectoderm. This revealed that differentiation of iPSCs to ectoderm revealed decreased protein O-GlcNAcylation and suggests that pathogenic differences in neural development occur at a subsequent neural progenitor stage, as ectoderm differentiates to neural progenitors.

### Neural progenitors have decreased O-GlcNAc levels

IPSC-derived neural progenitors cells (NPCs) form neural rosettes in cell culture models that mimic early stages of neural development, undergoing neurogenesis to give rise to neurons (and later glia) (<u>Pistollato et</u> <u>al., 2017</u>). To determine the effect of the OGT^skip^ variant on O-GlcNAc-mediated changes to ectodermal differentiation potential, NPCs were generated from the OGT^skip^ and OGT^corr^ IPSCs. Transcripts were assessed for the hallmarks of neural progenitors using a panel of common markers, resulting in no observable differences (Fig. 4A). Evaluation of protein levels in OGT^skip^ NPCs revealed significantly decreased OGT and OGA levels (to 45% and 41%, respectively) compared to OGT^corr^ NPCs (Fig. 4B,C). In undifferentiated IPSCs, decreased similarly OGT and OGA levels in OGT^skip^ lines did not affect total O-GlcNAc (Fig. 2A, B). Herein contrastwe observed a 50% decrease in total O-GlcNAcylation in OGT^skip^ NPCs. Further analysis of transcripts revealed no differences in either *OGT* and *OGA* between OGT^skip^ and OGT^corr^ NPCs (Fig. 4D). Therefore, there appears to be a shift in OGA regulation from transcriptional in iPSCs to entirely post-transcriptional in neural progenitors found in neural rosettes. Taken together, these data suggest that the differentiation of IPSCs to ectoderm and subsequently NPCs has resulted in a further decrease in O-GlcNAcylation.

**Figure 4:**
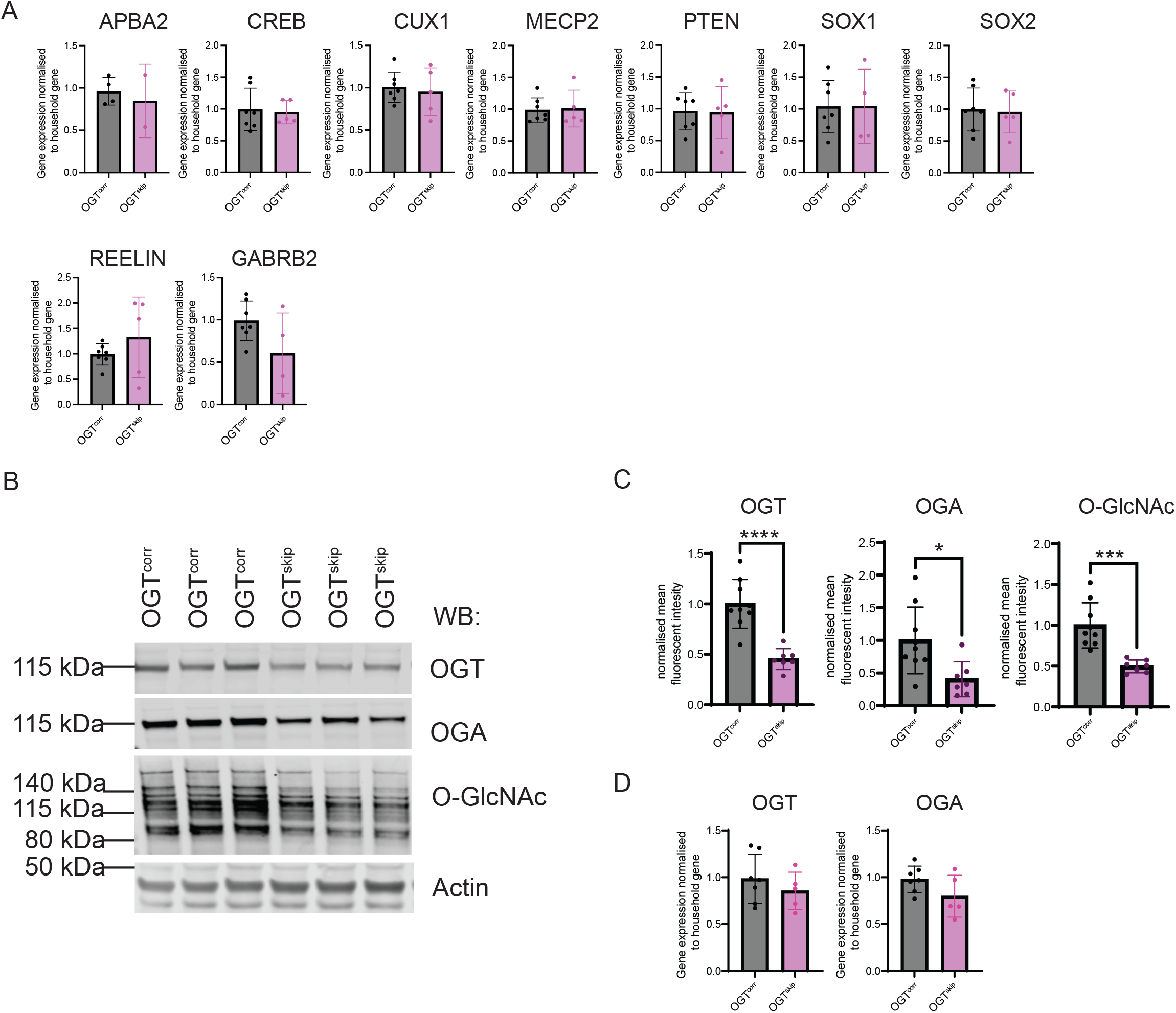
Neural progenitors and shifted O-GlcNAc regulation. **(a)** Neural progenitor cell specific markers evaluated by qPCR in the differentiated OGT^corr^ and OGT^skip^ cell lines. **(b)** Representative Western blot of neural progenitor cells and **(c)** Quantification of changes in OGT, OGA, and total O-GlcNAc. n = 3 experiments, 3 clonal lines/biological replicates for each. **(d)** Changes in transcription levels of OGT and OGA by qPCR.

### NPCs derived from OGTskip IPSCs are increased in size

Early in development, the neural tube gives rise to the central nervous system and forms multi-layered tube-like pseudostratified cells of defined orientation (<u>Paridaen and Huttner, 2014</u>). Differentiation of IPSCs to tissue with neural tube properties is not only reflected at molecular levels but also in terms of function and morphology (<u>Fedorova et al., 2019</u>, <u>Medelnik et al., 2018</u>, <u>Bino and Cajanek, 2023</u>). Several lines of evidence suggest that NPC structure may be affected by a range of cell signaling pathways (<u>Medelnik et al.,</u> <u>2018</u>). Because we observed differences in O-GlcNAcylation in NPCs, the morphology and cellular differences of neural progenitors were assessed. To investigate potential changes to NPC epithelial organization and apical lumen size, we examined morphology by immunofluorescence confocal microscopy (Fig. 5A, B). Distinctions in the overall morphology of the rosettes are visible, with fewer and larger rosettes apparent in OGT^skip^ patient cells. The apical lumens were significantly larger in the OGT^skip^ patient cells (median 20.3 μm) than OGT^corr^ variants (median 12.5 μm) (Fig. 5C), as measured by size of the rosette’s apical domain in ZO-1 stains.

**Figure 5:**
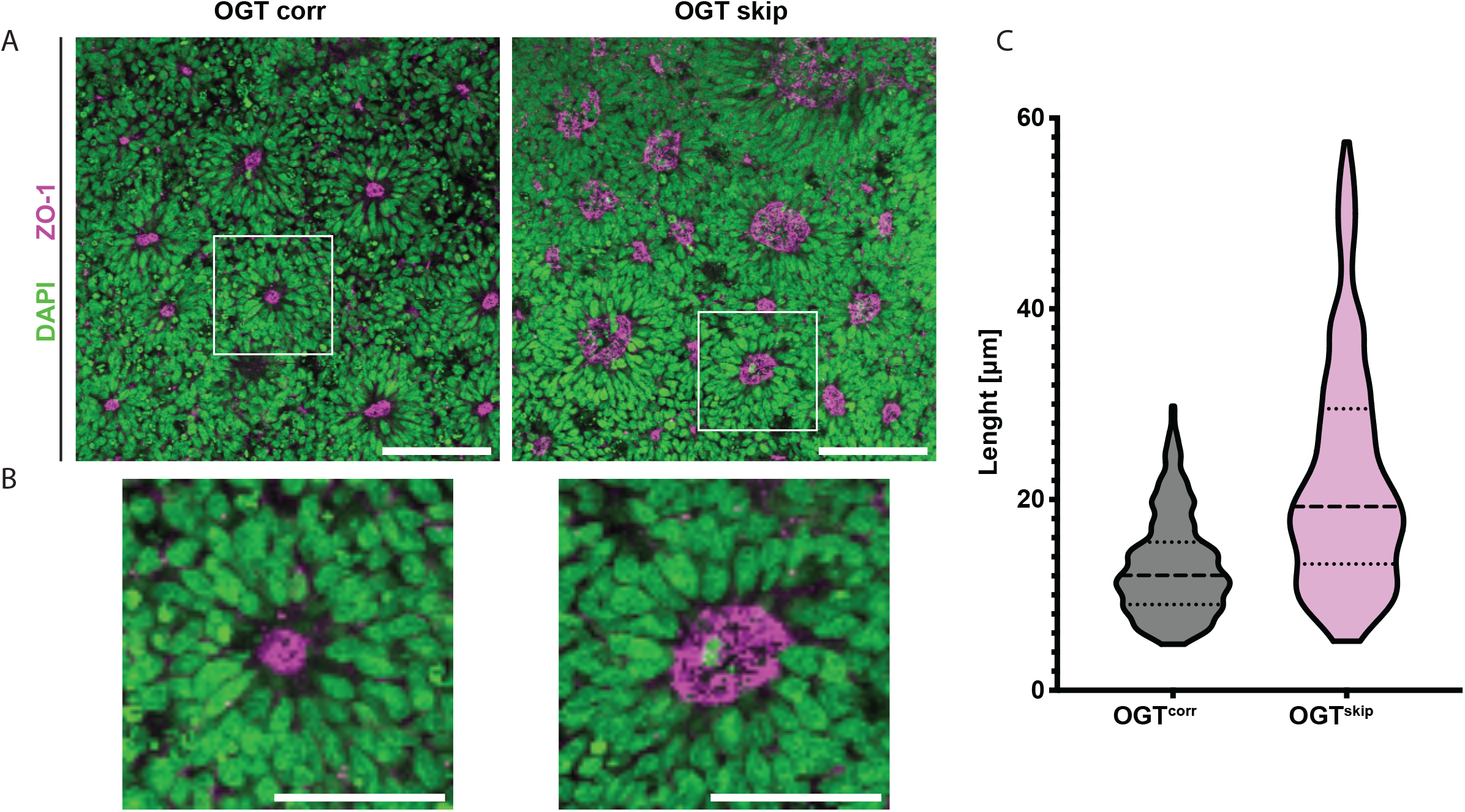
Apical size and morphological differences in neural progenitor cells. **(a)** Representative images of neural rosettes for OGT^corr^ and OGT^skip^ immunostained for DAPI (green) and ZO-1 (magenta). White box indicates insets. Scale bar, 100 μm. **(b)** Inset of single neural rosettes. Scale bar, 50 μm. **(c)** Quantification of apical membrane diameter, calculated across longest axis demonstrates non-normal distribution and increase in apical size in OGT^skip^ patient rosettes. Violin plot p value<0.0001 Mann Whitney test, dotted line represents median and two dashed lines represent two quartiles).

## Discussion

Several studies have revealed a critical role for O-GlcNAcylation in neurons. This modification directly supports the activity of neural synapse organizers (<u>Vosseller et al., 2006</u>, <u>Skorobogatko et al., 2011</u>, <u>Lee et</u> <u>al., 2021b</u>), and more specifically regulates a number of neuron-specific proteins resulting in suppression of neural excitability (<u>Lee et al., 2021b</u>). Moreover, catalytically-inactive OGA knock-in mice showed perinatal lethality (<u>Muha et al., 2021</u>, <u>O’Donnell et al., 2004</u>, <u>Yang et al., 2012</u>), while neuron-specific knockout of OGA resulted in changes of brain development (<u>Olivier-Van Stichelen et al., 2017</u>, <u>Su and Schwarz, 2017</u>). Neuronal loss of O-GlcNAcylation in OGT cKO mice leads to progressive neurodegeneration (<u>Wang et al.,</u> <u>2016</u>). While a neuroprotective role in neurodegeneration is proposed (<u>Wang et al., 2016</u>, <u>Wang et al., 2021</u>, <u>Cardozo et al., 2023</u>, <u>Muha et al., 2019</u>), the specific role of the O-GlcNAc modification in brain organization is less clear. Reports suggest both age- and disease-related changes in the brain upon O-GlcNAc pathway modulation (<u>Wheatley et al., 2019</u>, <u>Lee et al., 2021a</u>, <u>White et al., 2020</u>, <u>Lagerlof, 2018</u>). Hence, the O-GlcNAc pathway has a critical, temporal role in development, but this remains to be studied in the context of OGT-CDG brain development.

Advances in stem cell biology now enable the modeling of neurodevelopmental diseases at the earliest stages of phenotypic divergence, including for OGT-CDG (<u>Omelkova et al., 2023</u>, <u>Pravata et al., 2019</u>, <u>Selvan et al., 2018</u>). Despite significant progress, the neuropathological or developmental etiology of OGT-CDGs remains undetermined. Here, we have investigated the impact of a pathogenic OGT-CDG variant in IPSCs, the ectoderm, and NPCs, modelling each step of differentiation towards neural lineage. We find that IPSCs of the pathogenic OGT variant show significant decreases to their O-GlcNAc enzyme levels. Yet, these stem cells are remarkably robust to this variation, and able to maintain O-GlcNAc homeostasis, can self-renew, and retain pluripotency as demonstrated by their ability to differentiate into cell types representative of the three embryonic germ layers *in vitro*. However, we observed no increased differentiation of patient or genome-matched control cells, either in routine maintenance or directed differentiation.

Upon differentiation of IPSCs to ectoderm, which defines the germ layer responsible for the nervous system, significant decreases in total O-GlcNAc levels were apparent in the pathogenic OGT^skip^ variant. The functional importance of O-GlcNAcylation acquired upon differentiation suggests an increasing dependence of the pathway in differentiation. In particular, upon further differentiation of ectoderm to NPCs, the pathogenic variant maintained decreased levels of OGT, OGA, and O-GlcNAc suggesting that OGT and OGA levels may play roles in tissue function. This suppression of the O-GlcNAc pathway in OGT^skip^ cells results in phenotypic and morphological changes to NSC rosette structural organization. The changes observed in the neurodevelopmental stem cell model suggest the possibility of an early origin of OGT-CDG disability.

A balance of OGT and OGA activity is known to maintain O-GlcNAc homeostasis (<u>Decourcelle et al.,</u> <u>2020</u>, <u>Park et al., 2017</u>, <u>Zhang et al., 2014</u>). Previous studies have demonstrated a clear transcriptional regulation of OGA levels in the context of OGT-CDG associated mutations (<u>Vaidyanathan et al., 2017</u>, <u>Pravata et al., 2019</u>). Here, we observe different feedback mechanisms to regulate OGA. As IPSCs differentiate through ectoderm to NSCs, we observe a parallel compensatory transcriptional regulation of *OGA*, in line with previous evidence (<u>Decourcelle et al., 2020</u>). However, upon differentiating beyond NSCs, we observe a shift towards regulating OGA regulation at the post-transcriptional level. The precise mechanism for this shift is not clear, but may involve specific OGA to OGT feedback pathways (<u>Qian et al.,</u> <u>2018</u>, <u>Li et al., 2021</u>).

We examined neural development in the context of a pathogenic OGT^skip^ variant, finding robustness in stem cell function, but phenotypic defects arising in NSCs. Because this model recapitulates the organization of neural tube formation in physiology and neurological disease (<u>Curchoe et al., 2012</u>) (<u>Barak et al., 2022</u>), it is intriguing that changes in O-GlcNAc homeostasis are associated with increased NSC size and defective apical membrane organization. This result is consistent with decreased numbers of neurons upon OGT inhibition in human ESCs differentiated to neurons (<u>Andres et al., 2017</u>). Moreover, inhibition of OGA in mouse ESCs resulted in neural lineage differentiation defects (<u>Speakman et al., 2014</u>). Interestingly, suppression of OGT in mouse NSCs results in increased differentiation and decreased self-renewal (<u>Chen</u> <u>et al., 2021</u>). However, it is unclear if pathogenic OGT variants result in a similar decrease in OGT, or rather distinct O-GlcNAcomes. Neural progenitor size is tightly regulated (<u>Townshend et al., 2020</u>, <u>Bino and</u> <u>Cajanek, 2023</u>) and a differentiation defect at this neuroepithelium may link OGT-CDGs to a developmental imbalance. As NSCs form the basis for forebrain organoid development, it remains to be seen how the disorganization and size differences we observe for OGT^skip^ relate to organoid organization and function.

Taken together, this work suggests a critical role for O-GlcNAcylation in early neuroectoderm architecture, with robust compensatory mechanisms in the earliest stages of stem cell differentiation. The model system describe here provides a platform for dissecting the molecular mechanisms underpinning the neurodevelopmental defects that lead to OGT-CDG.

## Materials & Methods

### Generation of patient derived IPS cells

Primary fibroblasts from a patient carrying an exon4 skipping OGT-CDG mutation (<u>Willems et al., 2017</u>) were grown from skin biopsies, collected at the Radboud UMC expertise Center for Disorders of Glycosylation. Written informed consent for the re-use of fibroblasts was obtained from parents or legal guardians according to the approval of the local ethical committee (Radboud University Medical Center, the Netherlands; 2020-6588). Cell lines were cultured in DMEM/F12 (Life Technologies) supplemented with 10% FBS (Clone III, HyClones) and antibiotics streptomycin (100 μg/mL), penicillin (100 U/mL), and amphotericin (0.5 μg/mL) at 37 °C under 5% CO2.

Human induced pluripotent stem cells were derived, expanded and characterized by the Radboud UMC Stem Cell Technology Center. Fibroblasts were transduced at 80% confluence with four lentiviral vectors containing the pluripotency genes OCT3/4, NANOG, KLF4 and c-MYC. Transduced cells were incubated for 24 h at 37 °C, the lentiviruses were removed and cells washed three times with PBS. The following day, the transduced cells were transferred onto murine embryonic fibroblast (MEF)-coated plates and cultured for one month at 37 °C and 5% CO_2_ in a stem cell medium containing DMEM/Ham’s F-12 (cat. No. 1445C, Sigma Aldrich), 20% (v/v) Knock-out serum replacement (KOSR, cat. No. 10828-028, Thermo Fisher Scientific), MEM Non-essential amino acids (NEAA, cat. No. M7145, Sigma Aldrich), L-glutamine (cat. No. G8541, Sigma Aldrich), β-mercaptoethanol (cat. No. M3148 Sigma Aldrich), β-FGF (cat. No. F0291, Sigma Aldrich). One month after transduction, iPSC colonies were picked and expanded on Vitronectin-coated plates (cat. No. A14700, Thermo Fisher Scientific) in Essential E8 medium (cat. No. A1517001, Thermo Fisher Scientific).

### Chemical reagents

Unless otherwise stated, all chemical reagents were supplied by Sigma.

### Maintenance of cells

IPSCs were transferred to and maintained in TESR medium (<u>Ludwig et al., 2006</u>) supplemented with bFGF (30 ng/ml, Peprotech, Cat. No. 100-18B) and noggin (10 ng/ml, Peprotech, cat. No. 120-10C) on geltrex (Life technologies, Cat. No. A1413302). coated plates (10 μg/cm^2^) in a humidified atmosphere of 5% CO2 in air. Cells were routinely passaged twice a week using TrypLE Select (ThermoFisher, Cat. No. 12563011). For passaging TESR medium was further supplemented with 10 μM Rho kinase inhibitor Y-27632 (Tocris, Cat. No. 1254).

### CRISPR/Cas9 genome editing

All primers and DNA sequences were synthesized by IDT. hIPSCs were generated using CRISPR/Cas9 methodology using ssDNA as a repair template for correction of the mutation to reference genome. Transfection was performed using Neon electroporation system (ThermoFisher, MPK1025). The electroporation conditions used were 1150 V, 30 ms, and 1 pulse. Sequences of custom-made reagents are listed in Supplementary Table 1. Clonal selection was performed and screened using paired screening primers. PCR product was then digested with Fastdigest *Bve*I restriction enzyme (ThermoFisher, ER1741), which would not digest the PCR product if recombination was successful. Intact PCR products were confirmed by Sanger sequencing. Since the point variation was introduced in the intron leading to alternate splicing, we confirmed successful repair by extracting mRNA and performing one step RT-PCR (Takara Primescript High Fidelity, RR057B). Primer sequences and design are detailed in Supplementary Fig 1. The forward primer is located on exon 3 and reverse is located on exon 5, thus patient variant and corrected variant products will differ in size (275 bp vs 200 bp). The variation in splicing was additionally confirmed by Sanger sequencing.

### Differentiation protocols

For ectoderm differentiation, dual-SMAD inhibition (<u>Chambers et al., 2009</u>) as employed. Briefly, 60000 cells/cm^2^ were plated on dishes coated with 20 μg/cm^2^ and cultured in TESR medium supplemented with bFGF and ROCK inhibitor as described above (Day 0). The following day (Day 1), medium was changed to fresh mTESR with only bFGF. The following day (Day 2), medium was changed to hES base medium consisting of DMEM/F12 (ThermoFisher, 21331020) supplemented with 20% knockout replacement serum (ThermoFisher, 10828028), 1x GlutaMAX (ThermoFisher, 35050061), 1x NEAA (ThermoFisher, 11140050), 1x Pen/Strep (ThermoFisher, 15140122), and 0.1 mM β-mercaptoethanol (ThermoFisher, 31350010). This medium was further supplemented with 10 μm SB431542 (Tocris, cat no. 1614) and 100 nM LDN (Tocris, 6053). The following days (Day 3 and 4), medium was changed to fresh hES base supplemented with 10 μm SB431542 and 100 nM LDN. At day 6, medium changed to hES base and neurobasal culture medium at 3:1 ratio supplemented with 10 μm SB431542, 100 nM LDN. Neurobasal culture medium consists of neurobasal medium (Gibco, 21103049) supplemented with 1x N-2 supplement (ThermoFisher, 17502048), 1x B-27 supplement (ThermoFisher, 17504044), 1x GlutaMAX, 1x NEAA, and 1x Pen/Strep. At day 8, medium changed to hES base and neurobasal culture medium at 1:1 ratio supplemented with 10 μm SB431542 and 100 nM LDN. At day 10, medium was changed to hES base and neurobasal culture medium at 1:3 ratio supplemented with 10 μm SB431542 and 100 nM LDN. The ectoderm differentiated cells were examined or harvested at day 12.

### Immunoblotting

For immunoblotting, cells were washed once with PBS (Gibco, cat. No. 10010023) and lysed in appropriate amount of 1x Laemmli buffer (120 mM Tris-HCl (pH 6.8), 20% glycerol and 4% SDS). The samples were then sonicated (Bioruptor Twin, Diagenode). Protein concentration of the cell lysis was assessed with BCA protein assay (ThermoFisher, 23225) or Pierce™ 660 nm Protein Assay Reagent (ThermoFisher, 22660) supplemented with Ionic Detergent Compatibility Reagent (ThermoFisher, 22663). 20 μg of protein was boiled, loaded onto a 4-12% Bis-Tris gel (Invitrogen) and transferred on 0.2 μm nitrocellulose membrane (GE Healthcare cat.no 10600001). Following membrane blocking with 5% BSA in 1xTBS, antibodies were incubated overnight: Anti-O-GlcNAc Transferase (Abcam cat.no. 96718) Anti-O-GlcNAc (RL2, Novus cat. No NB300 1:1000), Anti-O-GlcNAcase (Sigma cat. No. HPA036141-100μl), Anti-Oct4 (Cell Signalling Technology cat. No.2890S), Anti-Sox2 (Cell Signalling Technology cat. No. 23064S), Anti-nanog (Cell Signalling Technology cat. No. 3580S) Anti-actin (Proteintech Cat. No. 66009-1-1g), anti GAPDH (Santa Crus cat. No. sc32233). The membranes were washed and then incubated with corresponding LI-COR secondary antibodies (1:20000). Signal was detected using LI-COR Odyssey scanning system and quantified using Image Studio Lite (LI-COR). Data were analyzed in Prism (GraphPad). Number of repeats per experiment is detailed in the figure legends.

### RT-qPCR

RNA was extracted from confluent cells using RNeasy kit (Qiagen cat. No. 74104). The quantification of the RNA concentration was measured (Nanodrop, Thermo) and mRNA was transcribed into cDNA (qScript cDNA Synthesis Kit, Quantabio). The qPCR reactions were set using 384-well plates in total volume of 10 μl of 10 ng cDNA, 250 nM primer mix and polymerase (PerfeCTa® SYBR® Green FastMix® for iQ™, Quanta or LUNA cat.no M3003E). qPCR was performed on BioRad CFX384 real time detection system (BioRad). The sequences of primers are supplied in supplementary table 1. The results are graphed using Prism (Graphpad). All qPCR experiments were repeated at least twice.

### Immunofluorescence

For immunofluorescence, cells were grown in μ-Slide 12-well (Ibidi, 81201). Standard procedure was used. Briefly, following washing cells were fixed with 4% formaldehyde, permeabilized with PBS-T, blocked in PBS-T supplemented with 5% BSA, followed by incubations with primary and secondary antibody. For immunofluorescence of the Sox2, Oct4 and Nanog transcription factors cells were fixed with 4% formaldehyde, and permeabilised in presence of ice-cold methanol. All other steps remained the same.

### Quantification of imaging

Following staining and imaging of neural rosettes, images were quantified manually using FIJI. Overall size of apical regions were determined by measuring the longest axis along the ZO-1 stained channel in rosettes, identified by morphology. Measurements were performed blinded.

### FACS

For the analysis of Sox2, Oct4 and nanog content, cells were washed and fixed with 4% PFA, then incubated with anti-nanog (Cell Signalling Technology cat. No. 3580S). Primary antibody was washed off and secondary antibody conjugated to Alexa 647 was used to visualise. Flow cytometry was performed on BD FACSCanto. Data were analysed using FlowJo 10 and Prism (GraphPad) software.

### Statistical analysis

Specific details of experimental, technical, and biological repeats are detailed in the Figure legends. All experiments were repeated at least 2 times with 3 biological repeats (clonal cell lines). For evaluation of Western Blots, statistical significance was determined by nested t-test over each biological repeat (clonal cell line). For all qPCR experiments and evaluation of microscopy, statistical significance was evaluated with Mann-Whitney test. Quantification of the light microscopy images was performed blinded.

## Acknowledgements

This work was funded by a Wellcome Trust Investigator Award (110061), a Novo Nordisk Foundation Laureate award (NNF21OC0065969) and a Villum Investigator Fonden award to D.v.A. We acknowledge support from the Human Induced Pluripotent Stem Cell, Light microscopy, and Flow Cytometry facilities in the School of Life Sciences, University of Dundee. The authors wish to thank the Radboud University Medical Centre Stem Cell Technology Center for generation of the iPSCs.

## Author contributions

M.M and D.v.A conceived the study; M.M performed experiments; L.D. advised and helped with cell culture A.T.F. designed primers and ssDNA; MM analysed data and MM and D.v.A. interpreted the data and wrote the manuscript.

## Supplementary figure legends

**Supplementary figure S1: Further quality control of generated IPSC stemness.**

**(a)** Stemness by qPCR of Yamanaka factors in the OGT^corr^ and OGT^skip^ cell lines.

**(b)** Representative confocal microscopy images of DAPI and Sox2 (as in Figure 1F),

**(c)** Oct4, and

**(d)** Nanog immunofluorescence at single cell resolution. Scale bar, 50 μm.

**(e)** Single-cell quantification of Yamanaka factors Sox2, Oct4 and Nanog by FACS. n=3.

**Supplementary Table 1:**
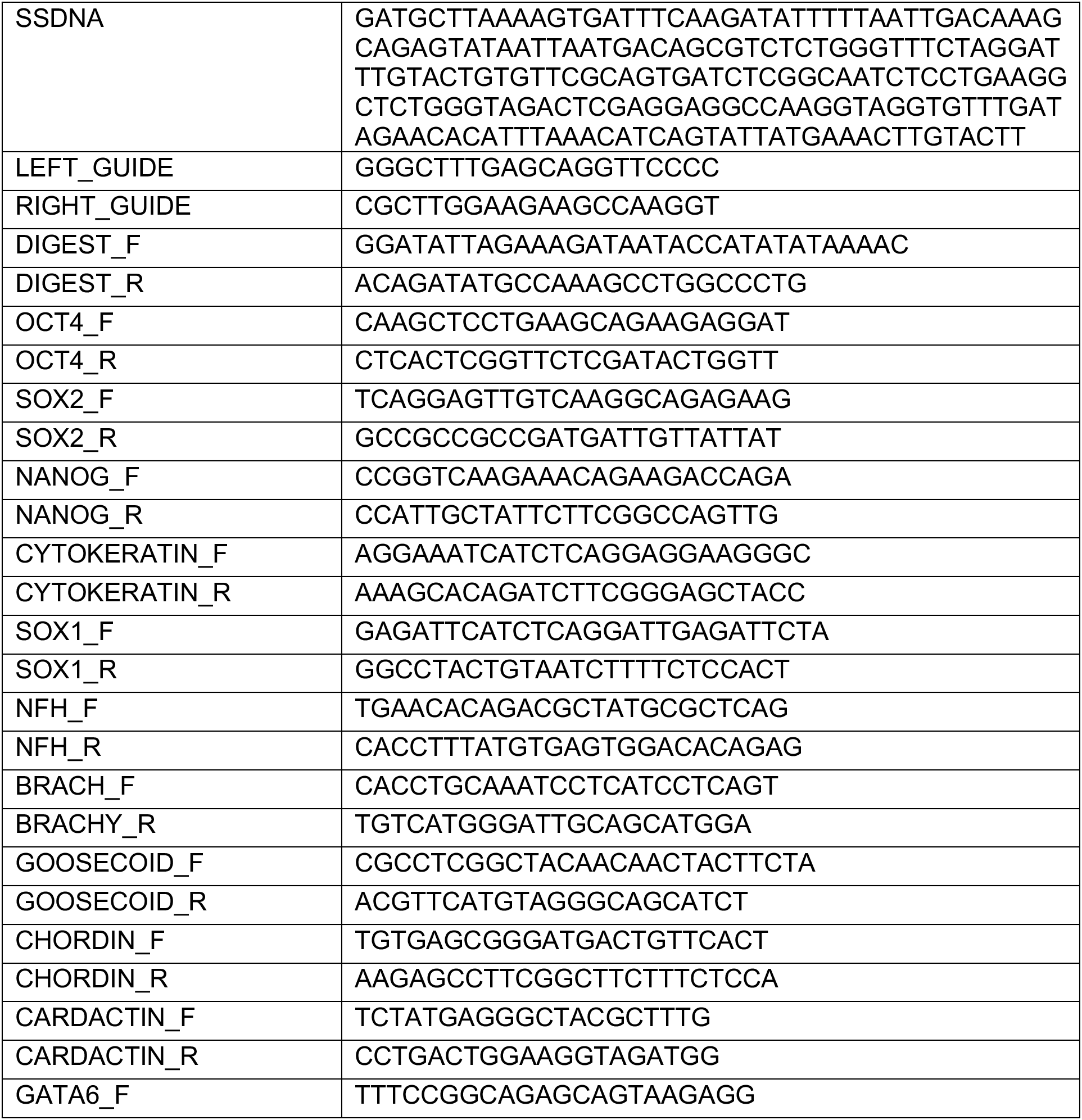
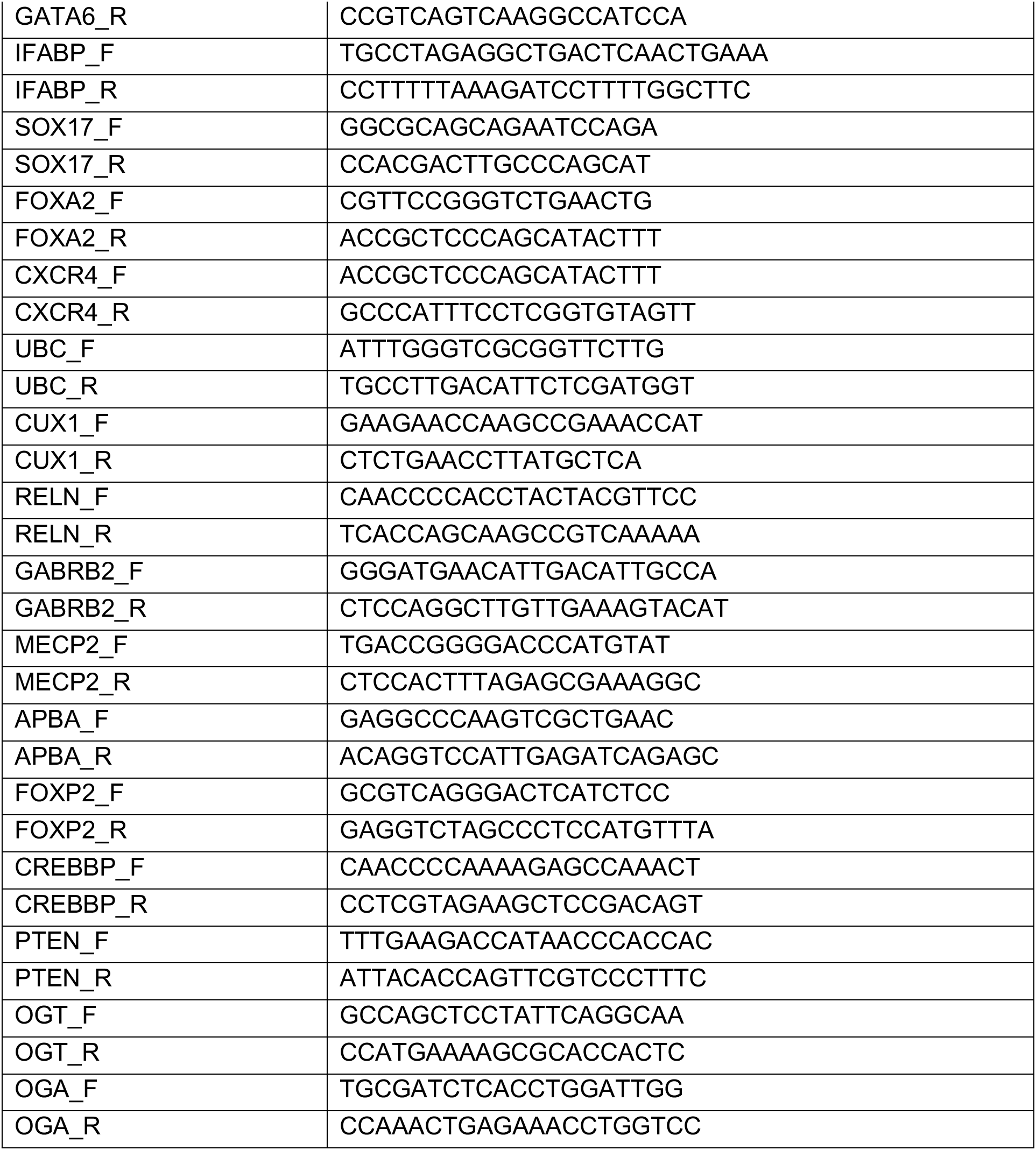
ssDNA, guides and primers used in this study.

